# Organelle specific protein profiling with light mediated proximal labeling in living cells

**DOI:** 10.1101/2019.12.28.890095

**Authors:** Zefan Li

**Affiliations:** College of Chemistry and Molecular Engineering, Peking University, Beijing, 100871, P. R. China

## Abstract

Organelle specific protein identification is essential for understanding how cell functions on a subcellular level. Here, we report a light mediated proximal labeling (LIMPLA) strategy for organelle specific protein profiling in living cells. In this strategy, various commercial mitochondria-localized fluorescent trackers, such as Mitoview 405 and Rhodamine 123, can activate 2-Propynylamine (PA) to label proximal proteins in mitochondria under illumination. PA tagged proteins are subsequently derivatized via click chemistry with azido fluorescent dye for imaging or with azido biotin for further enrichment and mass-spec identification. This strategy can be generalized to other organelles specific protein labeling. For example, proteins in nucleus are labeled by utilizing the commercial nucleus tracker DRAQ5. As compared with other chemical strategies for subcellular protein labeling, there are several advantages for this LIMPLA strategy. First, this approach allows minimal interference to the cell’s status by avoiding exogenous gene tansduction and some special treatment such as hydrogen peroxide or serum starvation. Second, all reagents used in this strategy are commercially available without additional synthesis work. Further, this strategy holds the potential for analyzing proximal proteins of specific macromolecules that can be tagged with fluorescent dye by metabolic labeling strategy.

## Introduction

Eukaryotic cells are elaborately subdivided into functionally distinct, membrane-enclosed compartments or organelles. Each organelle contains its own characteristic set of proteins, which underlies its characteristic structural and functional properties.^1^ For example, mitochondria, a double-membrane-bound organelle, are energy centers and plays crucial roles in energy conversion and calcium ions storage. Understanding how organelles function appropriately and specifically requires of proteomics in specific organelles.

Several protein proximity labeling (PL) methods combined with protein mass spectrometry have emerged for identification of organelle proteomics, in addition to traditional isolation of specific organelle method which may contain contamination and result in artifacts. The first PL method is based on a genetically encoded promiscuous labeling enzyme, which is targeted to a specific organelle and catalyzes covalent tagging of endogenous proteins proximal to the enzyme. BirA*, a mutant form of the biotin ligase enzyme BirA (BioID method)^2-4^ and APEX2, an engineered variant of soybean ascorbate peroxidase^5-7^ are mostly common used enzymes for this strategy. BirA* is capable of promiscuously biotinylating proximal proteins irrespective of whether these interact directly or indirectly with BirA*. This method has simple labeling protocol and need only biotin supplementation in order to initiate the labeling. Its disadvantages are long labeling time and background labeling signals due to endogenous biotin in living cells. The advantage of APEX2 is its short labeling time (only 1 min or less). However, the APEX2 method requires expression of exogenous proteins and the use of high concentration of H_2_O_2_, which is toxic to cells.

The second PL is based on antibody conjugated enzyme, such as horse radish peroxidase (HRP), which target to specific protein and tag endogenous proteins proximal to the HRP enzyme. The enzyme mediated activation of radical source (EMRS) method uses HRP conjugated antibody to targeted protein on cell membrane, where HRP mediates chemical reaction that converts aryl azide-biotin reagent to active radical species for proximal labeling.^8^ Biotin labeled proteins can then be high throughput analyzed. This method can only be used on cell membrane, however, due to membrane’s impermeability to antibodies. Selective proteomic proximity labeling using tyramide (SPPLAT) method^9^ is similar with EMRS and its substrate of HRP is changed to a cleavable biotin phenol. Biotinylation by antibody-recognition method^10^ ,in which fixed and permeabilized cells or tissues were treated with primary antibody for targeting protein of interest and then treated with a secondary HRP-conjugated antibody, H_2_O_2_ and phenol biotin for proximal protein biotinylation, extends EMRS to intra-cell,. However, this method need huge amounts of expensive antibody, is only applicable to fixed cells and may introduce artifacts due to long diffusion distance of biotin free radical.

Recently, mitochondria-localizable reactive molecules (MRMs)^11^ were designed to label and profile mitochondrial proteins, which eliminates the need for expression of exogenous proteins and H_2_O_2_ treatment. However, these reactive molecules may react with highly abundant proteins while in transit through the cytosol to mitochondria, leading to artifacts. Also, the MRMs labeling process requires starvation of serum for 6 h, which affects expression and phosphorylation of multiple proteins^12, 13^. The other potential drawback is that reagents in MRMs are not commercially available. This method also was applied to profiling protein in endoplasmic reticulum.^14^

Here, we describe a light mediated proximal labeling (LIMPLA) strategy for organelle specific protein profiling, which is independent of exogenous gene transfection and compatible with living cells. In the first version of this strategy, termed LIMPLA1 (Fig. 1), Alk-R123, an alkynyl analogue of rhodamine 123 (commercial mitochondrial fluorescent tracker, R123), was accumulated to mitochondria via membrane potential, and then upon illumination Alk-R123 labeled its proximal proteins in mitochondria via photochemical reaction. The labeled proteins bearing alkynyl functional group were reacted with azido fluorescent dye for imaging or azido-biotin for further enrichment and mass spectrometry analysis.

**Fig. 1.**
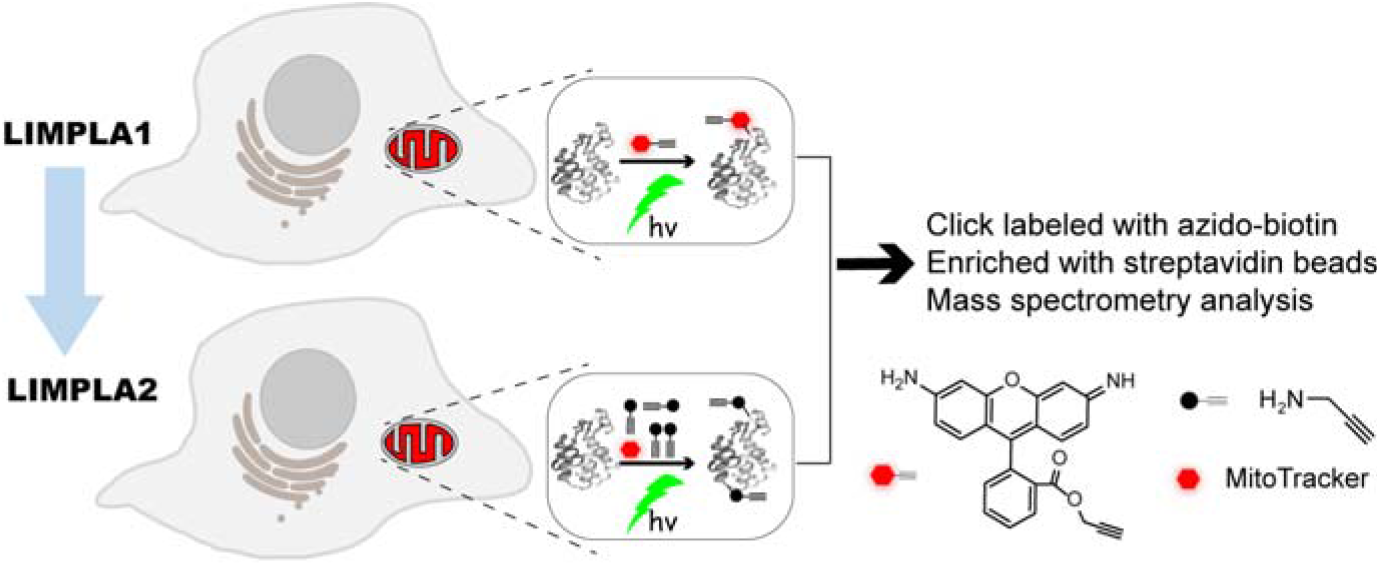
Schematic of light mediated proximal labeling (LIMPLA) strategy. LIMPLA1: Alk-R123, a derivative of mitochondria targeting dye Rhodamine 123, bearing alkyne group were accumulated in mitochondria, via mitochondria inner membrane potential. Upon illumination, these dyes can covalently label proximal proteins in situ. LIMPLA2: MitoTracker dye, such as Rhodamine123 or MitoTracker deep red, accumulated in mitochondria can mediate alkynyl substrate PA to label proximal proteins under illumination. The labeled proteins were reacted with azido dye for imaging or azido biotin for further enrichment and mass-spec identification.

## Results and discussions

### Development of LIMPLA1 for protein labeling in mitochondria

We began by synthesizing Alk-R123 and Az-R123 (an azido analogue of R123) (Scheme S1). To test if they can label bovine serum albumin (BSA) under illumination, these probes were incubated with BSA and illuminated with xenon light in vitro. Ith reaction with Azido-Cy5 or Alkynyl-Cy5, respectively, via Cu(I)-catalyzed azide-alkyne cycloaddition (CuAAC),^15, 16^ labeling signals on BSA were detected using in-gel fluorescence scanning. (Fig. S1). BSA treated with Alk-R123 or Az-R123 and light exhibit higher levels of signal than no light control group.

To validate cellular permeability and mitochondria targeting of Alk-R123, HeLa cells were incubated with Alk-R123 and Mito Tracker deep red, a commercial mitochondria marker. Confocal laser scanning microscopy (CLSM) showed that signals of Alk-R123 were highly co-localized with signals of Mitotracker deep red, which indicates that Alk-R123 selectively accumulated in mitochondria (Fig. S2). Mitochondrial localization was abolished by treatment with carbonyl cyanide m-chlorophenylhydrazone (CCCP),^17^ a standard uncoupler of the mitochondrial membrane potential (Fig. S2), which indicates the mitochondrial targeting of Alk-R123 was mediated with the high membrane potential of mitochondria.

To demonstrate the protein labeling ability in mitochondria with Alk-R123, HeLa cells stained with Alk-R123, were illuminated with xenon lamp. Cells were lysed and reacted with azido-Cy5 via CuAAC and then analyzed with in-gel fluorescence scanning. The results showed that HeLa cells treated with Alk-R123 and illumination group exhibit labeling signals (Fig. 2A lane4). As a control, HeLa cells only treated with Alk-R123 without illumination group exhibited level of labeling signals to the levels of background (Fig. 2A lane2). Labeling signals were almost abolished by CCCP (Fig. 2A lane3), as the membrane potential of mitochondria disappeared and Alk-R123 were not accumulated in mitochondria. Furthermore, the fluorescence imaging results showed that green labeling signals of HeLa cells were well correlated with red anti-HSP60 (HSP60: a mitochondrial proteins) signals (Fig. 2B), which indicates the labeling specifically occurred in mitochondria. These results were in accordance with the results of in gel fluorescence scanning, which indicates that the Alk-R123 accumulated in mitochondria can label proteins in situ with illumination. The Alk-R123 labeling signals were incubation time- and illumination time-dependent (Fig. S3-4). Similarly, Az-R123 can also be used for protein labeling in mitochondria as shown in Fig. S5.

**Fig. 2.**
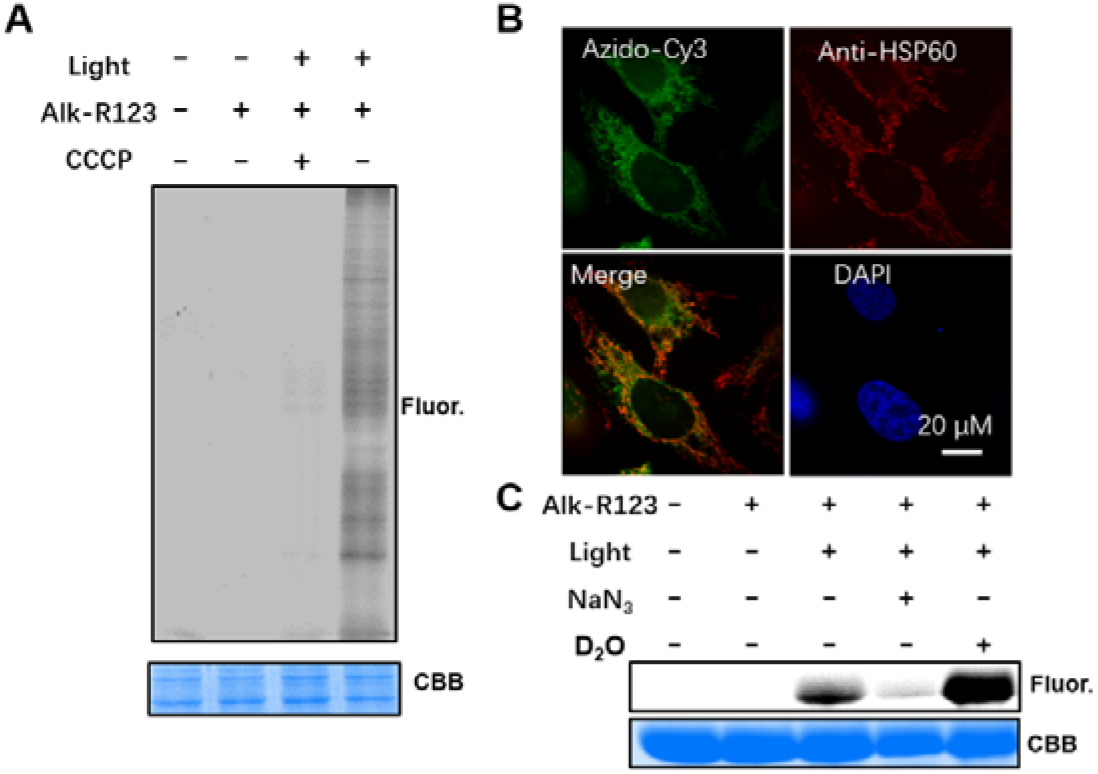
Mitochondrial protein labeling with LIMPLA. (A) HeLa Cells were stained with 10 μM Alk-R123 for 2h. Cells were then illuminated for 5 min. Cell lysate were reacted with azido-Cy5 via CuAAC for in gel fluorescence scanning. (B) After treatment of Alk-R123 and illumination, cells were fixed, permeabilized and clicked with azio-Cy3. Cells were immunostained with Anti-HSP60. Green channel denotes azido-Cy3 signal. Red channel denotes HSP60 signal. Blue channel denotes DAPI signal. (C) 50 μM Alk-R123, 2 mg/mL BSA and 1mM PA (or + 5 mM NaN3) in H2O or D2O were illuminated for 2 min. The labeled proteins were reacted with azido-Cy5 for in gel fluorescence analysis.

It was speculated that the light mediated labeling was catalyzed by singlet oxygen generated in the process of dye photobleaching.^18^ The sodium azide, a singlet oxygen quencher^19^ ,can reduce the BSA labeling signals, while D_2_O which increases singlet oxygen lifetime, can enhance the BSA labeling signals (Fig. 2C). Taken together, these data suggest that the singlet oxygen plays an important role in the photo-labeling.

The LIMPLA1 strategy is related to cell labelling via photobleaching (CLaP),^20, 21^ a method designed to specifically tagging individual cells based on photobleaching. In CLaP, a laser is used to crosslink fluorescent biotin conjugates to cell membrane of living cells via free radicals generated by photobleaching. Instead of cell membrane labeling, our method utilized crosslink of proteins in mitochondria via photo reaction.

### Development of LIMPLA2 for protein labeling in mitochondria

In order to further increase the photo-labeling signals, we began by increasing the concentration of Alk-R123. But when the concentration of Alk-R123 was increased above 20 μM, Alk-R123 bound nonspecifically with non-mitochondrial proteins which resulted in background signals. We speculated that Alk-R123 can be split to two parts—one is R123 for mitochondrial targeting and generating singlet oxygen upon illumination and the other is alkynyl substrate for protein labeling catalyzed by singlet oxygen (**Fig. 1**). This may increase the protein labeling efficiency because the concentration of labeling reagent alkynyl substrate can be dramatically boosted to mM level without impairing the mitochondrial targeting. To test the idea, we selected N-(4-aminophenethyl) pent-4-ynamide (AY) bearing benzenamine structure similar with rhodamine and a simple primary amine PA as the alkynyl substrates (chemical structure showed in **Fig. 3D**). In this test, HeLa cells were stained with 10 μM R123, respectively, incubated with 5 mM AY or PA, and then illuminated with xenon lamp. Cell lysate were reacted with azido Cy5 via CuAAC and analyzed with in gel fluorescent scanning. As shown **in Fig. 3A**, labeling signals in PA+ R123 group are stronger than AA+ R123 group. Besides AY substrate, we also tested (o, m, p)-alkynyl benzenamine as the alkynyl substrate in HeLa cell for mitochondrial protein labeling with R123 and under illumination. All the three benzenamine substrates can show certain signals and PA still shows the highest signals among all tested substrates (Fig. S6). Combining these results together, we select the PA as the alkynyl substrate for the following experiment. To compare with the Alk-R123, labelling signals in R123+ PA group are much higher (**Fig. 3B**). In **Fig. S7**, labeling signals in CCCP group were dramatically impaired and signals in no light or no R123 group were completely disappeared, which is consistent with the results in LIMPLA1. After treated with R123, PA and illumination, cells were in situ reacted with azido Cy3, and the labeling signal were well correlated with immunofluorescence signals of a mitochondrial protein HSP60 (**Fig. 3C**), which indicated that PA labeled proteins were located in mitochondria. We termed this new version of <u>l</u>ight <u>m</u>ediated <u>p</u>roximal <u>la</u>beling as LIMPLA2. Compared with LIMPLA1, LIMPLA2 has much higher labeling efficiency and has no need to do any organic synthesis, as compounds used in LIMPLA2 are all commercial available.

**Fig. 3.**
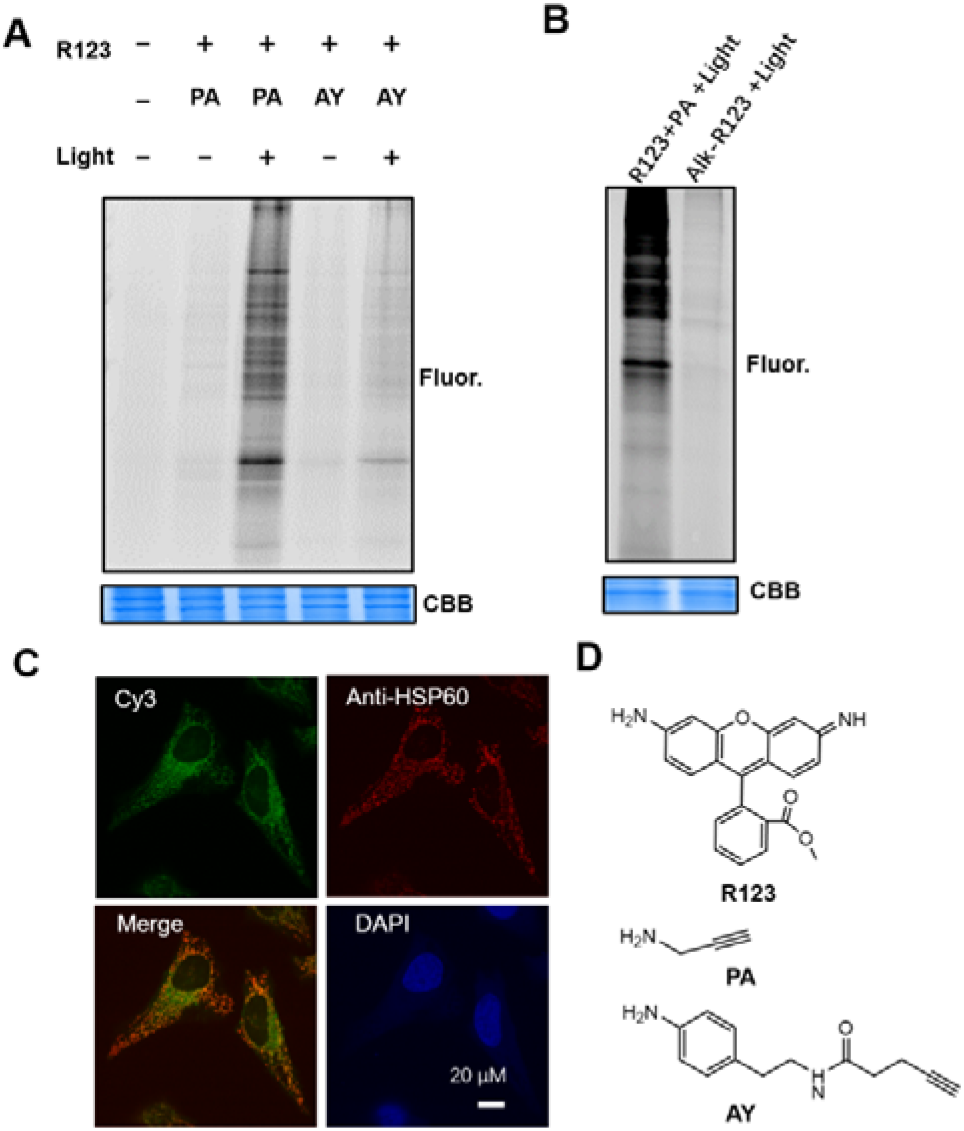
Mitochondrial protein labeled with LIMPLA2. (A) In gel fluorescence analysis of R123+PA and R123+AY labeling. HeLa Cells were stained with 10 μM R123 for 3h. Cells were treated with 5 mM AY or PA for extra 5 min. Cells were then illuminated for 5 min. Cell lysate were reacted with azido Cy5 for in gel fluorescence scanning. (B) In gel fluorescence analysis of R123+PA and Alk-R123 labeling. (C) After treatment with R123, PA and illumination, cells were fixed, permeabilized and clicked with azio-Cy3. Cells were immunostained with Anti-HSP60. Green channel denotes azido-Cy3 signal. Red channel denotes HSP60 signal. Blue channel denotes DAPI signal. (D) Chemical structures of R123, PA and AY.

We next asked whether LIMPLA2 can be used for RNA labeling. HeLa cells were treated with R123 for 2h (negative group were treated with CCCP), cultured with 5 mM PA for 5min and then illuminated with xenon lamp for 5min. RNA were isolated from these cells and click labeled with azido biotin. The labeled RNA were analyzed by dot blot analysis. R123 treated groups showed significant labeling signals, while the signals were abolished by CCCP which shows that the labeled RNA should be located in mitochondria (Fig. S8). This result indicated that the LIMPLA2 strategy can be extended to RNA labeling.

This LIMPLA2 strategy may be applied to be analyze proximal proteins of other macromolecules, such as glycan, combined with metabolic labeling strategy. For example, similar with a reported method termed “protein oxidation of sialic acid environments” (POSE) for mapping the proximal protein of sialic acids in situ^22^, unnatural sialic acid bearing alkynyl handle metabolically installed on cell membrane can be click labeled with azido fluorescent dye and then sialic acid’s proximal proteins can be labeled with LIMPLA2 by utilizing the glycan conjugated fluorescent dye.

LIMPLA2 is inspired by the method for RNA and protein labeling mediated by fluorophore eosin,^23-26^ in which eosin is spatially confined in subcellular region via conjugation with halo tag protein or other subcellular targeting moiety, and then eosin will mediate PA for labeling proximal macromolecule upon illumination. However, eosin derivatives in this method are not commercially available and lengthy chemical synthesis work is unavoidable; eosin may be relatively toxic to cultured cell even under natural light compared with common dye as eosin is always used as a photosensitizer and has high singlet oxygen quantum yield,^27, 28^ which also means eosin may have higher photo-labeling efficiency. When this manuscript is under preparation, an excellent method for mapping spatial RNA is published.^29^ In this paper, the authors utilize a singlet oxygen generator protein miniSOG as photosensitizer which is expressed in mitochondria and can mediate RNA labeling with PA under illumination.

### Improvement of LIMPLA2 for mitochondrial protein labeling

We wonder if there is another mitochondrial tracker which can further increase the LIMPLA2’s labeling efficiency. Three new mitochondrial trackers mitoView 405 (mito405), Tetramethylrhodamine (TMRM), MitoTracker deep red were tested. In this experiment, HeLa cells were, respectively, stained with various mitochondria trackers, and then treated with PA and light. In this situation, the cell lysates were reacted with azido-biotin instead of azido-Cy5 or other fluorescent dye in order to avoid the spectrum overlap between azido bearing dye and various MitoTrackers. The labeling results were analyzed by biotin western blot. As shown in **Fig. 4A**, all of three new trackers groups exhibit labeling signals and mito405 had the strongest labeling signals among all groups. Then the labeling signals of mito405 in mitochondria were detected by fluorescent imaging. The imaging results show that these signals were well correlated with immunofluorescence signals of a mitochondrial protein HSP60, which indicated that labeled proteins were located inside mitochondria (**Fig. 4B**). In **Fig. 4C**, labeling signals in CCCP group were impaired and labeling signals in no light or R123 group were completely abolished. The labeling intensity is positively correlated with concentration of mito405 (**Fig. S9**) and PA (**Fig. S10**). We applied LIMPLA2 to label mitochondrial proteins in primary neuron culture. Cortical neuron cells were stained with mito405, incubated with PA and illuminated for xenon light. The labeling results were analyzed with in situ fluorescent imaging and in gel fluorescence scanning (**Fig. S11**), which shows that LIMPLA2 can be used for labeling mitochondrial proteins in primary neurons.

**Fig. 4.**
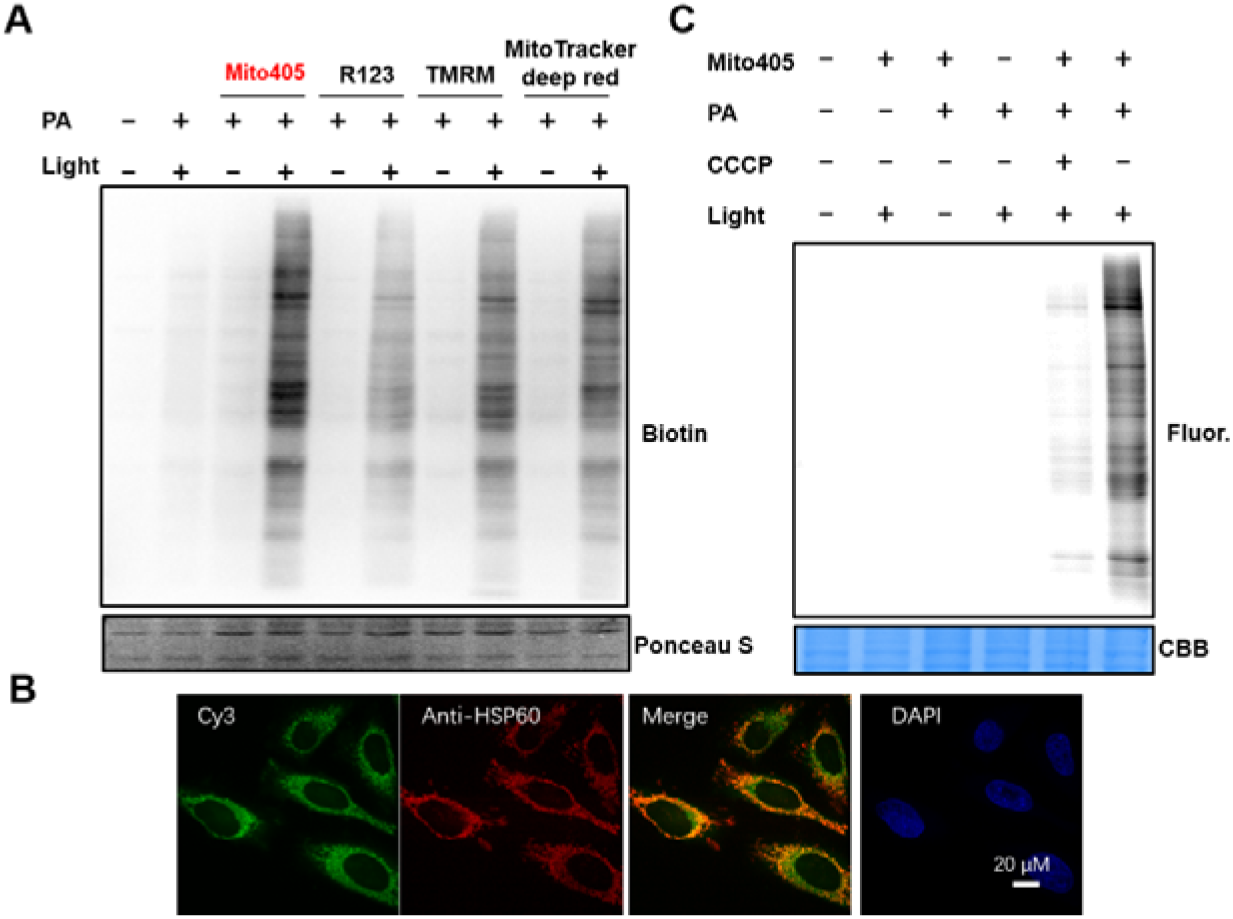
Evaluation of various MitoTrackers in LIMPLA2 (A) HeLa Cells were, respectively, stained with 10 μM R123 or 0.5μM other MitoTrackers for 1h. Cells were treated with 5 mM PA for 5 min and then illuminated for 5 min. Cell lysate were reacted with azido-biotin for in western blot analysis. (B) After treatment with mito405, PA and illumination, cells were fixed, permeabilized and clicked with azido Cy3. Cells were immunostained with Anti-HSP60. Green channel denotes azido Cy3 signal. Red channel denotes HSP60 signal. Blue channel denotes DAPI signal. (C) In gel fluorescence scanning analysis of protein labeling with mito405 based LIMPLA2.

Further, we verified that far red light can mediate LIMPLA2 labeling. It was reported that far red light with much longer wavelengths can deeply penetrate tissue and decreased photo-toxicity compared with blue or green light^30^. We chose far red dye Mitotracker deep red (EM/EX spectrum shown in Fig. S12) for testing *in vitro*. BSA mixed with Mitotracker deep red dye and PA were illuminated with xenon lamp or xenon lamp with various wavelength light filter. After protein precipitation for removing PA in the system, BSA were click labeled with azido Cy3 dye. In gel fluorescence scanning showed that far red group with 600-660nm band pass filter or 650 nm-long pass filter can show significantly stronger signals than negative group, but the signals were weaker than 400 nm-long pass filter or full length (xenon lamp only without filter) group (Fig. S12). These results demonstrated far red light should be compatible with LIMPLA2.

### Proteomic analysis of labeled protein by LIMPLA2

We utilized stable isotope dimethyl labeling based quantitative mass spectrometry to identify the labeled proteins by LIMPLA2. HeLa cells treated with mito405, PA and illumination and cells treated with PA and illumination are, respectively, as experimental and negative control group (**Fig. 5A**). After treatment, HeLa cells were lysed, reacted with azido-biotin. Biotin labeled proteins were pulled down with streptavidin beads and were digested with trypsin overnight and subsequently labeled with heavy formaldehyde ^13^CD_2_O for experimental group and light formaldehyde CH_2_O for the control group. Finally, samples in experimental group and control group were mixed in a 1: 1 ratio (in volume) and subjected to LC-MS/MS for identification and quantification. We did a biological repeat by swapping CH_2_O and ^13^CD_2_O labeling. We set a heavy/ light or light /heavy ratio of 5.0 as the cutoff to exclude any non-specific proteins. In replicate 1 and 2, we have identified 283 and 200 proteins, respectively. About 60% proteins are mitochondrial localizing proteins and the ratio is much higher than the mitochondrial protein ratio in human proteome (**Fig. 5C**). These non-mitochondrial proteins are likely resulting from misdistribution of the trackers mito405 and mischaracterization of the labeled peptide in the LC-MS/MS analysis.^11^ Combining the identified protein in 2 replicates, we identified 329 proteins and 188 mitochondria localizing proteins (**Fig. 5D and 5E**).

**Fig. 5.**
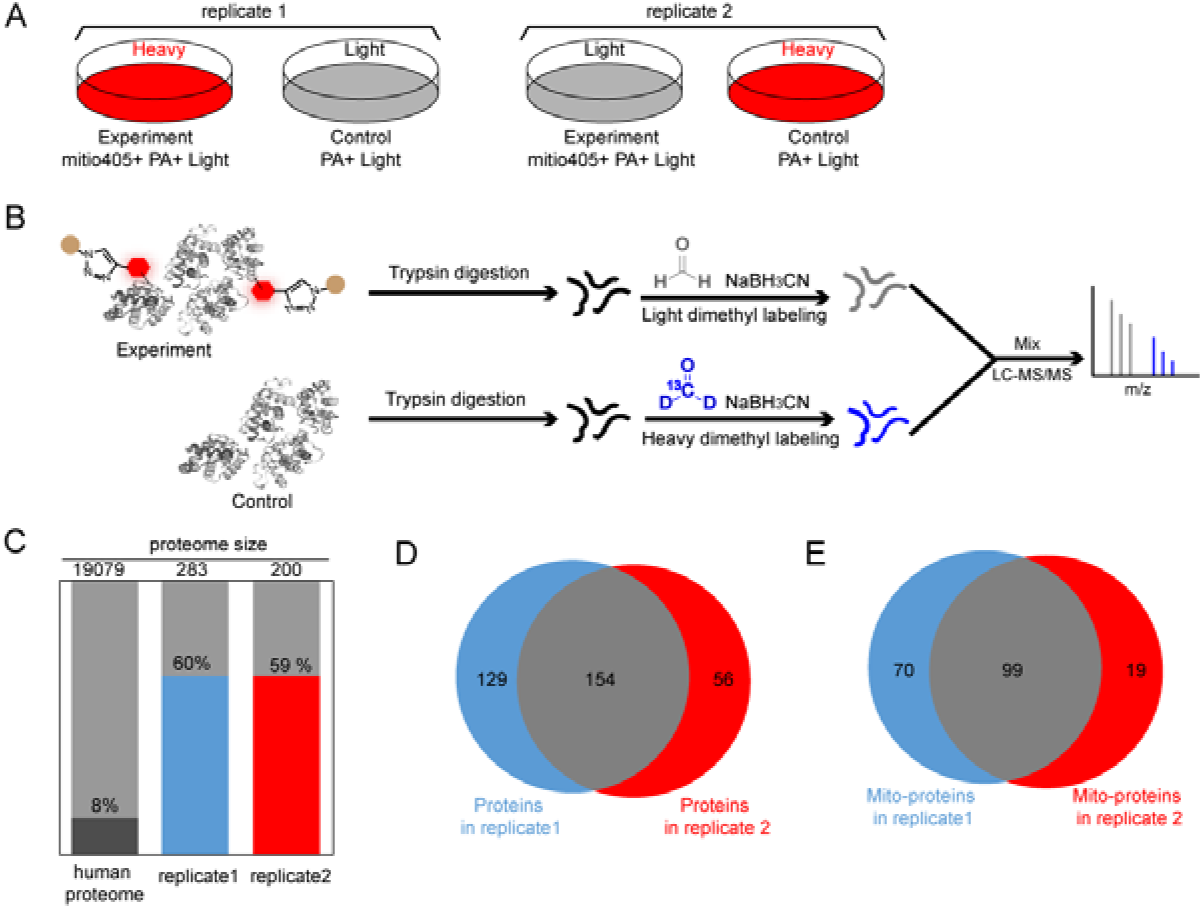
Identification of proteins labeled with LIMPLA2. (A) Experimental setup. HeLa cells treated with mito405, PA and illumination and cells treated with PA and illumination are, respectively, as experimental and negative control group. In replicate 1, peptides from experimental group reacted with heavy formaldehyde; peptides from control group reacted with light formaldehyde. In replicate 2, peptides from experimental group were reacted with light formaldehyde; peptides from control group reacted with heavy formaldehyde. (B) The schematic of sample preparation for dimethyl labeling based quantitative mass spectrometry. (C) Mitochondrial specificity analysis of identified proteins by using the protein ontology information from the UniProt database. (D) Overlap of identified proteins. (E) Overlap of identified mitochondria locating proteins.

### Application of LIMPLA2 for protein labeling in nucleus

We tested if LIMPLA2 can be extended to label proteins in other organelles. We selected DRAQ5, a cell nucleus tracker, as it was reported that DRAQ5 can be photo excited to generate reactive oxygen species that catalyze DAB polymerization on chromatin in the nucleus for visualizing 3D chromatin structure. HeLa cells were stained with DRAQ5, treated with PA and then illuminated with xenon light. Cell lysates were reacted with azido Cy5 via CuAAC and the in gel fluorescence scanning results show that DRAQ5 can mediate the LIMPLA2 labeling (**Fig. 6A**). Imaging results shows that click signals were well correlated with DRAQ5 signals (**Fig. 6B**), which indicated that the proteins labeling were specifically confined in cell nucleus. In addition to mitochondria and nucleus, LIMPLA strategy is supposed to be useful in labeling and profiling of other organelles, such as endoplasmic reticulum.

**Fig. 6.**
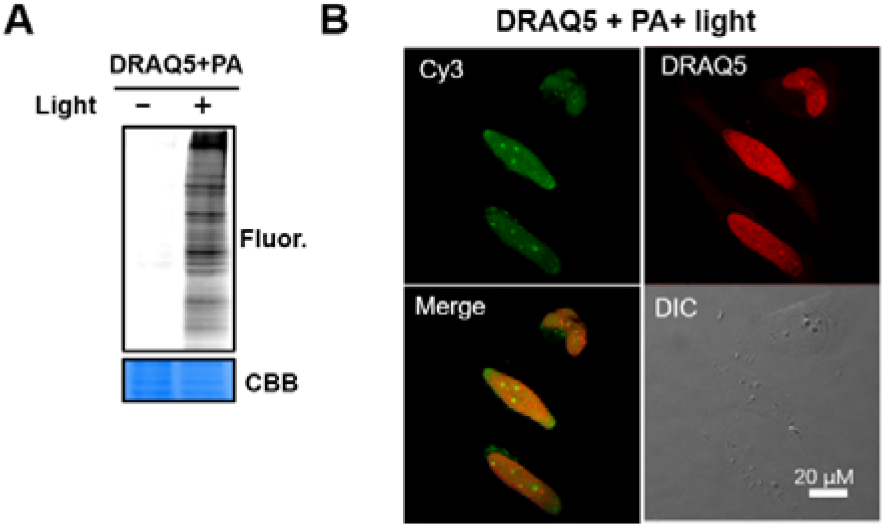
Proteins in nucleus labeled with LIMPLA2. (A) HeLa Cells were stained with 10 μM DRAQ5 for 2h. Cells were treated with 5 mM PA for 5 min and then illuminated for 5 min. Cell lysate were reacted with azido-Cy3 for in gel fluorescence scanning. (B) After treatment with DRAQ5, PA and illumination, cells were fixed, permeabilized and clicked with azido-Cy3 via CuAAC. Green channel denotes azido-Cy3 signal. Red channel denotes DRAQ5 signal.

## Experimental section

### Reagents and materials

Alkynyl-Cy5, azido-Cy5, azido-Cy3, biotin-PEG3-azide, THPTA and BTTAA were purchased from Click Chemistry Tools (Scottsdale, AZ, USA). 2-Propynylamine (PA), 2-ethenylbenzenamine, 3-ethenylbenzenamine, 4-ethenylbenzenamine, Rhodamine 123 (R123), and Tetramethylrhodamine (TMRM) were purchased from J & K Scientific (Beijing, P. R. China). MitoTracker deep red, TRIzol reagent, Pierce streptavidin agarose resins and DRAQ5 (5 mM solution) were purchased from Thermol Fisher scientific (Waltham, MA USA). MitoView 405 (mito405) was purchased from Biotium Inc. (Fremont, CA, USA). Xenon light source (300W) was purchased from Bochengguangdian (Nanjing, P. R. China); its emission spectrum ranges from 300-1000nm. Formaldehyde-13C, D2 solution was purchased from Sigma-Aldrich (St. Louis, MO, USA)

### Chemical Synthesis of Alk-R123 and Az-R123

Rhodamine 110 (30 mg, 82 μmol), trimethylamine (TEA, 24.4 mg, 164 μmol), dimethylaminopyridine (DMAP, 1mg, 82 μmol) and propynol (46 mg, 820 μmol) or 2-azidoethan-1-ol (71.5 mg, 820 μmol) were dissolved in 1 mL dichloromethane in a 5 mL tube on ice. When the mixed solution recovered to RT, EDCI (31 mg, 164 μmol) was added. Then the tube was placed on shaker for overnight reaction. After centrifugation of the tube, the supernatant was evaporated and products was purified by flash chromatography to red solid. Alk-R123 (10.1 mg, 30.6%) and Az-R123 (12.3 mg, 32.3%) were obtained.

### Labeling and imaging of cells

HeLa cells were cultured in DMEM supplemented with 10% FBS. Cortical neuron cells were cultured in DMEM supplemented with 10% FBS, 200 μg/mL geneticin. HeLa cells were seeded into 24 well plate and were grown to 90% confluence. In CCCP group, HeLa cells were pretreated with 10 μM CCCP for 5min. HeLa cells were stained with an indicated mitochondrial dye (0.5 μM mito405, 0.5 μM TMRM, 0.5 μM MitoTracker deep red or 10 μM AR123) for 1 h. Cells were washed with PBS for 3 times. If the dye is mito405, TMRM, MitoTracker or R123 deep red, cells were incubated with 5 mM PA or other alkyne compounds for 5 min and then cells were washed with PBS for 3 times. Cells were then illuminated with xenon light for 5 min.

After illumination, cells were rinsed three times with PBS, fixed with 3.7% paraformaldehyde solution for 10 min, and rinsed three times with PBS. Cells were then permeabilized with 0.1% Triton-X100 in PBS for 10 min, and rinsed for three times with PBS, followed by adding of PBS containing 50 μM Cy5 azide, 2.5 mM sodium ascorbate, and BTTAA-CuSO_4_ complex (50μM CuSO_4_, BTTAA: CuSO in a 2:1 molar ratio) at rt.^32, 33^ After reaction, cells were washed four times with 1% Tween 20 in PBS and once with PBS. Cells were incubated with anti-HSP60 antibody Rabbit (Invitrogen), followed by AF647-conjugated anti-rabbit secondary antibody (Invitrogen). Cells were then stained with Hoechst 33342 prior to imaging. Fluorescence imaging experiments were conducted on an inverted fluorescence microscope (Nikon-TiE) equipped with a 40X 1.3 NA oil immersion objective lens, five laser lines (Coherent OBIS, 405 nm, 488 nm, 532 nm, 561 nm, 637 nm), a spinning disk confocal unit (Yokogawa CSUX1), and two scientific CMOS cameras (Hamamatsu ORCA-Flash 4.0 v2).

### In-gel fluorescence scanning of labeled protein

After illumination, cells were lysed with RIPA lysis buffer (1% Nonidet P 40, 1% sodium deoxycholate, 0.1% SDS, 50 mM triethanolamine pH 7.4, 150 mM NaCl) containing protease inhibitor cocktail (ETDA-free, CWBIO). Cells were lysed by pipetting and agitation. Protein concentrations were determined using the BCA Protein Assay Kit (Pierce, USA) and normalized to 1 mg/ mL with lysis buffer. Samples were then reacted with 100 µM Cy5 azide in a 60 µL reaction containing premixed BTTAA-CuSO_4_ complex (50 µM CuSO_4_, BTTAA: CuSO_4_ in a 2:1 molar ratio) and 2.5 mM freshly prepared sodium ascorbate. The samples were vortexed on an IKA shaker for 1 h at rt. Reacted samples were resolved on 10% SDS-PAGE gels. The gel was incubated in destaining solution (50% methanol, 40% H_2_O, 10% glacial acetic acid) for 5 min followed by H_2_O for an additional 5 min prior to scanning. The gel was scanned on a Typhoon FLA 9500 laser scanner (GE Healthcare, USA).

### Dot blot analysis of labeled RNA

HeLa cells were treated with 10 or 50 μM R123 (or supplemented with 10 μM CCCP). After 1h, cells were treated with 5mM PA and then illuminated for 5 min. RNA were isolated from cells per the manufacturer’s instructions and were click labeled with azido-biotin via CuAAC in a 40 uL reaction containing 100 μM biotin-PEG3-azide, 0.5 mM CuSO4, 2 mM THPTA and 5 mM sodium ascorbate for 30 min at RT with shaking. Then RNA were purified with RNA clean & concentrator kit. Equal amount of purified RNA was loaded on Immobilon-Ny⍰+⍰membrane (Merck Millipore, INYC00010-1) and crosslinked to the membrane by 350 nm UV light from an ultraviolet crosslinker (Analytik Jena). The membrane was blocked with 5% BSA at room temperature for 1⍰h and incubated with Streptavidin-HRP (Pierce, 21124) at RT for 1⍰h. The membrane was washed three times with TBST (50⍰mM Tris, 150⍰mM NaCl, 0.1% Tween-20, pH⍰7.4–7.6) for 10⍰min each time, incubated in Clarity Western ECL Substrate (Bio-Rad, 1705061) and then imaged on a ChemiDoc imaging system (Bio-Rad).

### Affinity enrichment of labeled proteins

Labeled cells were lysed and proteins were reacted with biotin-PEG3-azide by CuAAC for 1 h at rt. Unreacted biotin-PEG3-azide was removed by the addition of ice-cold methanol, placed at -80□ for 1 h to precipitate proteins. After washed with ice-cold methanol twice, proteins were redissolved in 1.2% SDS in PBS buffer by sonication. The biotinylated proteins were then captured by incubation with streptavidin-agarose resins (Pierce) for 3 h at rt. The resins were extensively washed sequentially with 2% SDS in PBS, 8M urea, 2.5 M sodium chloride in PBS, PBS for three times and water for three times.

### Stable isotope dimethyl labeling of peptides

After enrichment using biotin-PEG3-azide, the resins were reduced with 5 mM dithiothreitol (DTT) for 15 min, and alkylated with 10 mM Iodoacetamide (IAA) for 30 min in 100 mM TEBA buffer. After centrifugation and washing to remove the remaining DTT and IAA, 500 ng/μL trypsin was added to the resin and the sample was incubated for 16 h at 37l □ with agitation. The samples were dried by vacuum centrifugation, reconstitute in 100 mM TEAB, and 4% (v/v) CH_2_O or ^13^CD_2_ O were added for light and heavy dimethyl labeling respectively, followed by the addition of 0.6 M NaBH3CN. After incubation with agitation for 30 min at room temperature, the labeling reaction was quenched by adding 1% (v/v) ammonia solution for 10 min. Then formic acid was added for further quenching the reactions. Light and heavy samples were mixed in a 1:1 ratio, dried by vacuum centrifugation and reconstituted in water. The peptides samples were fractionated with a fast sequencing (Fast-seq) workflow using dual reverse phase (RP) high performance liquid chromatography (HPLC). The first dimension of high pH RP chromatography was performed on an Agilent 1260 infinity quaternary LC by using a durashell RP column (5 μm, 150 Å, 250 mm × 4.6 mm i.d., Agela). Mobile phases A (2% acetonitrile, adjusted pH to 10.0 using NH_3_·H_2_O) and B (98% acetonitrile, adjusted pH to 10.0 using NH_3_·H_2_O) were used to develop a gradient. The solvent gradient was set as follows: 5–8% B, 2 min; 8–18% B, 11 min; 18–32% B, 9 min; 32–95% B, 1 min; 95% B, 1 min; 95–5% B, 2 min. The tryptic peptides were separated at an eluent flow rate of 1.5 ml/min and monitored by UV at 214 nm. The temperature of column oven was set as 45°C. Eluent was collected every minute. The samples were dried under vacuum.

For LC-MS/MS analysis, the samples were reconstituted in 0.2 % formic acid, loaded onto a 100 μm x 2 cm pre-column and separated on a 75 μm×15 cm capillary column with a laser-pulled sprayer. Both columns were packed in-house with 4 μm C18 bulk material (InnosepBio, P. R. China). An Easy nLC 1000 system (Thermo Scientific, USA) was used to deliver the following HPLC gradient: 5-35% B in 75 min, 35-75% B in 10 min, then held at 75% B for 15 min (A = 0.1% formic acid in water, B = 0.1% formic acid in acetonitrile). The eluted peptides were sprayed into an Orbitrap XL mass spectrometer (Thermo Scientific, USA) equipped with a nano-ESI source. The mass spectrometer was operated in data-dependent mode with a full MS scan (375-1600 m/z) in FT mode at a resolution of 30000 followed by CID (Collision Induced Dissociation) MS/MS scans on the 15 most abundant ions in the initial MS scan.

### Identification and quantification of proteins

The data were analyzed using MaxQuant software (v. 1.5.3.30).^34^ Spectra extracted from the raw file were searched against Uniprot Homo sapiens database and contaminants database. Digestion enzyme was specified as trypsin with up to two missed cleavages. Variable modifications included methionine oxidation (+15.9949), N-terminal protein acetylation (+42.0106), and a fixed modification of cysteine carbamidomethylation (+57.0215). The light label was set as DimethLys0 (C2H4) and the heavy label was set as DimethLys6 (Cx2Hx4). Precursor mass tolerance was less than 4.5 ppm after mass recalibration by MaxQuant. Fragment ion tolerance was 0.5 Da. Protein and peptide false discovery rates were fixed at 1% and estimated using a decoy database search performed by MaxQuant.

## Conclusions

In summary, we have developed a strategy LIMPLA for organelle protein profiling in living cells. Various commercially available mitochondria fluorescent trackers can be used for mitochondrial protein labeling, which was validated by fluorescence imaging, in gel fluorescence and Mass spectrometry. Besides protein, mitochondrial RNA can be labeled using this strategy. According to BSA labeling experiment with Mitotracker deep red, the strategy is compatible with far red light, which may extend the usage of the strategy to animal tissue. By combining with quantitative mass-spec technique, this strategy allowed the identification of mitochondrial proteins. Further, this strategy was generalized to labeling protein in nucleus by utilizing commercial nucleus fluorescent tracker DRAQ5.

## Acknowledgments

We thank F. Yuan for preparing the cortical neuron cells, Y. Li for offering the AY compound and P. Lv for helping performing mass spectrometry experiment. Z. Li was supported in part by the Postdoctoral Fellowship of Peking-Tsinghua Center for Life Sciences.

Scheme S1 Chemical synthesis of Alk-R123 and Az-R123.

Fig. S1 BSA labeling with Alk-R123 and Az-R123 under illumination. 50 μM Alk-R123, Az-R123 and 2 mg/mL BSA in H2O were illuminated for 2 min. Then, cold methanol was added for protein precipitation, placed at -80 ºC overnight. After centrifugation, the supernatant was discarded, and the protein pellets were wash for 2 times with cold methanol. The proteins pellets were resuspended with 0.1% SDS buffer and were reacted with azido-Cy5 or alkyne-Cy5 using CuAAC.

Fig. S2 Mitochondria targeting of Alk-R123. HeLa cells were incubated with 0.1 μM Mitotracker deep red, 10 μM AR 123 and 10 μM CCCP for 1h, and then were imaged by confocal laser scanning microscopy (CLSM).

Fig. S3 Illumination time dependence of Alk-R123 labeling. HeLa cells were incubated with 10 μM Alk-R123 for 2 h, cells were illuminated with light for 0, 1, 5, 10, 15 min. Cell lysates were reacted with azido-Cy5, and then were analyzed by in gel fluorescence.

Fig. S4 Incubation time dependence of Alk-R123 labeling. HeLa cells were incubated with 10 μM Alk-R123 for 0, 15m, 30m, 1h, 2h, 3h, cells were illuminated with light for 5min. Cell lysates were reacted with azido-Cy5, and then were analyzed by in gel fluorescence.

Fig. S5 Az-R123 labeling mitochondria protein. (A) Imaging analysis of HeLa cells labeled with Az-R123. Cells were treated with 10μM Az-R123 for 2 h, and then illuminated with xenon lamp for 5 min. After fixation and permeabilization, cells were reacted with azide-Cy5 via CuAAC. Red channel denotes azide-Cy5 signal. Blue channel denotes DAPI signal. (B) In gel fluorescence analysis of HeLa cells labeled with Az-R123. Cells were treated with 10μM Az-R123 for 2 h, and then illuminated with xenon lamp for 5 min. Cell lysate were reacted with azido-Cy5 for in gel fluorescence.

Fig. S6 Labeling efficiency of PA and o-, p-, m-aminophenylacetylenes mediated by R123 and light. HeLa Cells were stained with 10 μM R123 for 3h. Cells were treated with 5 mM AY or PA for extra 5 min. Cells were then illuminated for 5 min. Cell lysate were reacted with azido Cy5 for in gel fluorescence analysis.

Fig. S7 In gel fluorescence analysis of mitochondrial proteins labeling with R123 and PA based LIMPLA2.

Fig. S8 Dot blot analysis of RNA labeling by LIMPLA2. HeLa cells were treated with 10 or 50 μM R123 (or supplemented 10 μM CCCP) for 2 h, cultured with 5 mM PA for 5 min. RNA were isolated from these cells and click labeled with azido biotin. Purified RNA were analyzed by dot blot analysis.

Fig. S9 PA concentration dependent labeling mediated by mito405 and PA based LIMPLA2. Fig. S10 Mito405 concentration dependent labeling mediated by mito405 and PA based LIMPLA2.

Fig. S11 Cortical neuron cells labeling with LIMPLA2. (A) Cortical neuron cells were stained with 0.1 μM mito405 for 2h, incubated with 5 mM PA for 5 min and illuminated for 5 min. Cells were clicked labeled with azido-Cy3 and analyzed with in situ fluorescent imaging (B) In gel fluorescence scanning analysis of labeling.

Fig. S12 BSA labeling mediated by Mito tracker deep red with various wavelengths of light. (A) Excitation and emission spectrum of Mitotracker deep red. (B) In gel fluorescence scanning analysis of protein labeling mediated with various wavelengths of light. 1mg/mL BSA mixed with 1 μM Mitotracker deep red and 1mM PA were illuminated with xenon lamp or xenon lamp with 400nm-, 540-590nm, 600-660nm filter for 2min. After protein precipitation, BSA were click labeled with azido Cy3.

## References

1. J. R. Yates, 3rd, A. Gilchrist, K. E. Howell and J. J. Bergeron, Nat. Rev. Mol. Cell Biol., 2005, 6, 702–714.

2. R. Varnaite and S. A. MacNeill, Proteomics, 2016, 16, 2503–2518.

3. K. J. Roux, D. I. Kim, M. Raida and B. Burke, J. Cell Biol., 2012, 196, 801–810.

4. T. C. Branon, J. A. Bosch, A. D. Sanchez, N. D. Udeshi, T. Svinkina, S. A. Carr, J. L. Feldman, N. Perrimon and A. Y. Ting, Nat. Biotechnol., 2018, 36, 880–887.

5. H.-W. Rhee, P. Zou, N. D. Udeshi, J. D. Martell, V. K. Mootha, S. A. Carr and A.Y. Ting, Science, 2013, 339, 1328–1331.

6. S. A. Myers, J. Wright, R. Peckner, B. T. Kalish, F. Zhang and S. A. Carr, Nat. Methods, 2018, 15, 437–439.

7. S. Markmiller, S. Soltanieh, K. L. Server, R. Mak, W. H. Jin, M. Y. Fang, E. C. Luo, F. Krach, D. J. Yang, A. Sen, A. Fulzele, J. M. Wozniak, D. J. Gonzalez, M. W. Kankel, F. B. Gao, E. J. Bennett, E. Lecuyer and G. W. Yeo, Cell, 2018, 172, 590–604.

8. N. Kotani, J. Gu, T. Isaji, K. Udaka, N. Taniguchi and K. Honke, Proc. Natl. Acad. Sci. U.S.A., 2008, 105, 7405–7409.

9. X.-W. Li, J. S. Rees, P. Xue, H. Zhang, S. W. Hamaia, B. Sanderson, P. E. Funk, R. W. Farndale, K. S. Lilley, S. Perrett and A. P. Jackson, J. Biol. Chem., 2014, 289, 14434–14447.

10. D. Z. Bar, K. Atkatsh, U. Tavarez, M. R. Erdos, Y. Gruenbaum and F. S. Collins, Nat. Methods, 2017, 15, 127–133.

11. Y. Yasueda, T. Tamura, A. Fujisawa, K. Kuwata, S. Tsukiji, S. Kiyonaka and I. Hamachi, J. Am. Chem. Soc., 2016, 138, 7592–7602.

12. V. A. Levin, S. C. Panchabhai, L. Shen, S. M. Kornblau, Y. Qiu and K. A. Baggerly, J. Proteome Res., 2010, 9, 179–191.

13. S. Cooper, FASEB J., 2003, 17, 333–340.

14. A. Fujisawa, T. Tamura, Y. Yasueda, K. Kuwata and I. Hamachi, J. Am. Chem. Soc., 2018, 140, 17060–17070.

15. V. V. Rostovtsev, L. G. Green, V. V. Fokin and K. B. Sharpless, Angew. Chem., Int. Ed., 2002, 41, 2596–2599.

16. C. Besanceney-Webler, H. Jiang, T. Zheng, L. Feng, D. Soriano Del Amo, W. Wang, L. M. Klivansky, F. L. Marlow, Y. Liu and P. Wu, Angew. Chem. Int. Ed. Engl., 2011, 50, 8051–8056.

17. W. H. Gao, Y. M. Pu, K. Q. Luo and D. C. Chang, J. Cell. Sci., 2001, 114, 2855–2862.

18. J. R. Lepock, J. E. Thompson and J. Kruuv, Biochem. Biophys. Res. Commun., 1978, 85, 344–350.

19. M. Bancirova, Luminescence, 2011, 26, 685–688.

20. L. Binan, J. Mazzaferri, K. Choquet, L. E. Lorenzo, Y. C. Wang, B. Affar el, Y. De Koninck, J. Ragoussis, C. L. Kleinman and S. Costantino, Nat. Commun., 2016, 7, 11636.

21. L. Binan, F. Belanger, M. Uriarte, J. F. Lemay, J. C. Pelletier De Koninck, J. Roy, E. B. Affar, E. Drobetsky, H. Wurtele and S. Costantino, elife, 2019, 8, e45239.

22. Q. Li, Y. Xie, G. Xu and C. B. Lebrilla, Chem. Sci., 2019, 10, 6199–6209.

23. L. Li, J. Liang, H. Luo, K. M. Tam, E. C. M. Tse and Y. Li, Chem. Commun. (Camb.), 2019, 55, 12340–12343.

24. Y. Li, M. B. Aggarwal, K. Ke, K. Nguyen and R. C. Spitale, Biochemistry, 2018, 57, 1577–1581.

25. Y. Li, M. B. Aggarwal, K. Nguyen, K. Ke and R. C. Spitale, ACS Chem. Biol., 2017, 12, 2709–2714.

26. Y. Li, K. Ke and R. C. Spitale, Biochemistry, 2019, 58, 379–386.

27. T. J. Deerinck, M. E. Martone, V. Lev-Ram, D. P. Green, R. Y. Tsien, D. L. Spector, S. Huang and M. H. Ellisman, Journal of Cell Biology, 1994, 126, 901–910.

28. J. T. Ngo, S. R. Adams, T. J. Deerinck, D. Boassa, F. Rodriguez-Rivera, S. F. Palida, C. R. Bertozzi, M. H. Ellisman and R. Y. Tsien, Nat. Chem. Biol., 2016, 12, 459–465.

29. P. Wang, W. Tang, Z. Li, Z. Zou, Y. Zhou, R. Li, T. Xiong, J. Wang and P. Zou, Nat. Chem. Biol., 2019, 15, 1110–1119.

30. J. Icha, M. Weber, J. C. Waters and C. Norden, Bioessays, 2017, 39, 1700003.

31. H. D. Ou, S. Phan, T. J. Deerinck, A. Thor, M. H. Ellisman and C.C. O’Shea, Science, 2017, 357, eaag0025.

32. Z. Li, Y. Zhu, Y. Sun, K. Qin, W. Liu, W. Zhou and X. Chen, ACS Chem. Biol., 2016, 11, 3273–3277.

33. Y. Li, W. Liu, Q. Tang, X. Fan, Y. Hao, L. Gao, Z. Li, B. Cheng and X. Chen, ACS Chem. Biol., 2019, 14, 182–185.

34. J. Cox and M. Mann, Nat. Biotechnol., 2008, 26, 1367–1372.

