## Supplemental file 1 for "Organelle specific protein profiling with light mediated proximal labeling in living cells"

Zefan Li*

College of Chemistry and Molecular Engineering, Peking University, Beijing, 100871, P. R. China

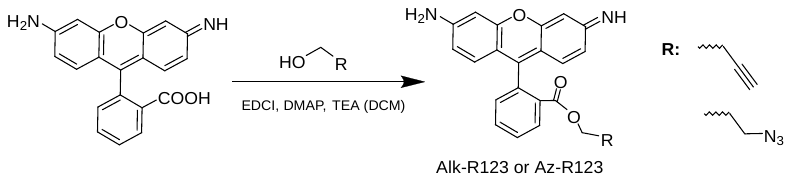

Scheme S1	Chemical synthesis of Alk-R123 and Az-R123.

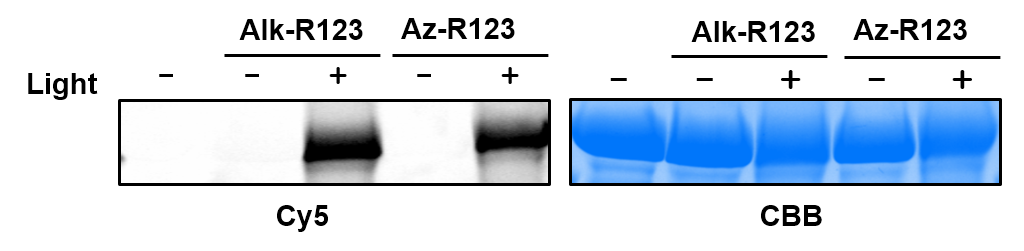

Fig. S1	BSA labeling with Alk-R123 and Az-R123 under illumination. 50 μM Alk-R123, Az-R123 and 2 mg/mL BSA in H_2_O were illuminated for 2 min. Then, cold methanol was added for protein precipitation, placed at -80 ℃ overnight. After centrifugation, the supernatant was discarded, and the protein pellets were wash for 2 times with cold methanol. The proteins pellets were resuspended with 0.1% SDS buffer and were reacted with azido-Cy5 or alkyne-Cy5 using CuAAC.

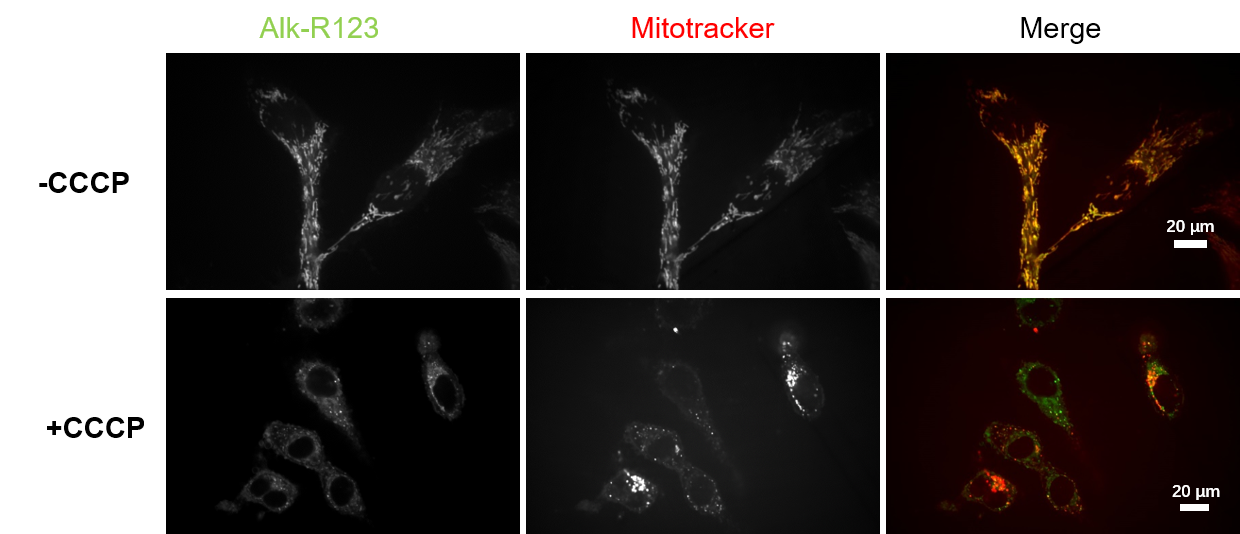

Fig. S2	Mitochondria targeting of Alk-R123. HeLa cells were incubated with 0.1 μM Mitotracker deep red, 10 μM AR 123 and 10 μM CCCP for 1h, and then were imaged by confocal laser scanning microscopy (CLSM).

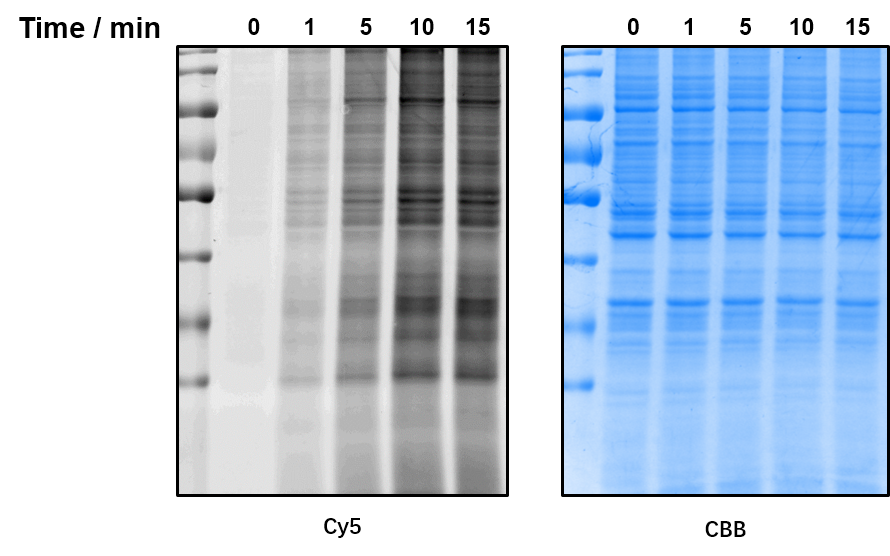

Fig. S3	Illumination time dependence of Alk-R123 labeling. HeLa cells were incubated with 10 μM Alk-R123 for 2 h, cells were illuminated with light for 0, 1, 5, 10, 15 min. Cell lysates were reacted with azido-Cy5, and then were analyzed by in gel fluorescence.

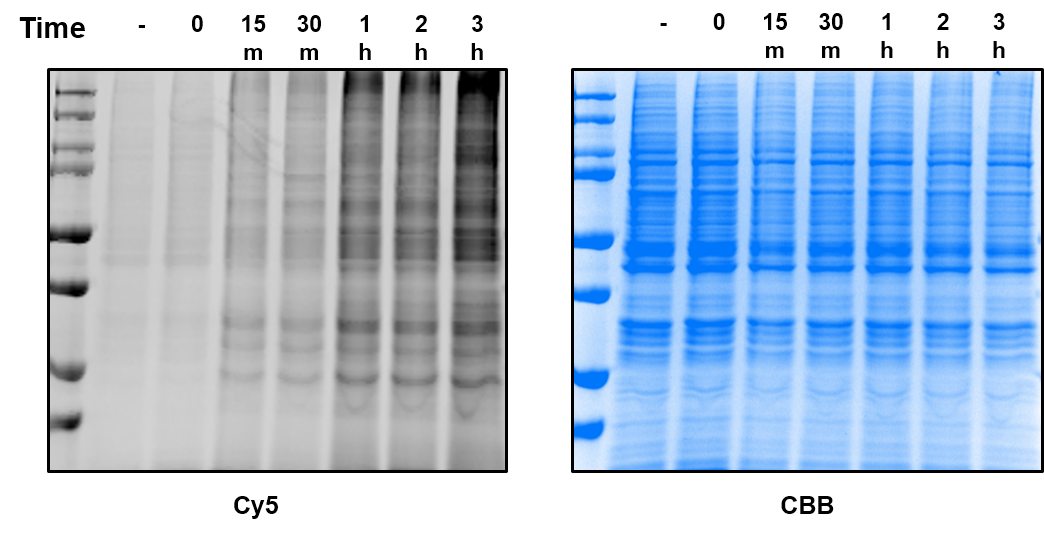

Fig. S4	Incubation time dependence of Alk-R123 labeling. HeLa cells were incubated with 10 μM Alk-R123 for 0, 15m, 30m, 1h, 2h, 3h, cells were illuminated with light for 5min. Cell lysates were reacted with azido-Cy5, and then were analyzed by in gel fluorescence.

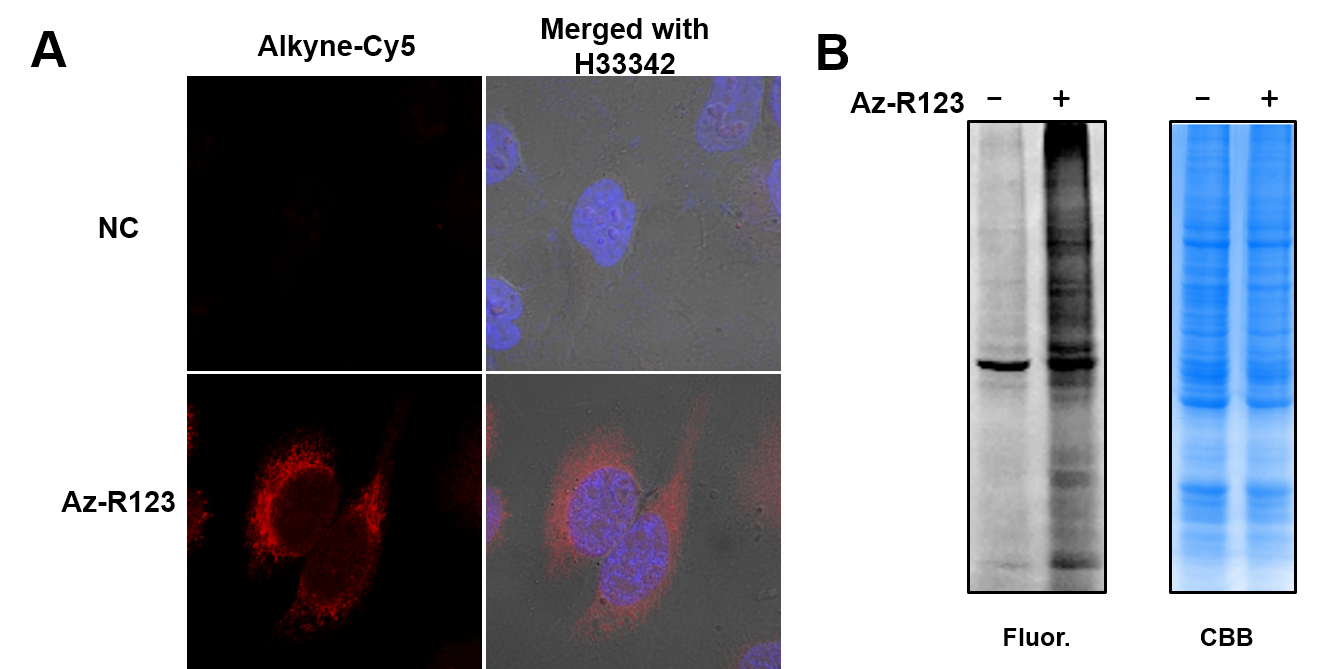

Fig. S5	Az-R123 labeling mitochondria protein. (A) Imaging analysis of HeLa cells labeled with Az-R123. Cells were treated with 10μM Az-R123 for 2 h, and then illuminated with xenon lamp for 5 min. After fixation and permeabilization, cells were reacted with azide-Cy5 via CuAAC. Red channel denotes azide-Cy5 signal. Blue channel denotes DAPI signal. (B) In gel fluorescence analysis of HeLa cells labeled with Az-R123. Cells were treated with 10μM Az-R123 for 2 h, and then illuminated with xenon lamp for 5 min. Cell lysate were reacted with azido-Cy5 for in gel fluorescence.

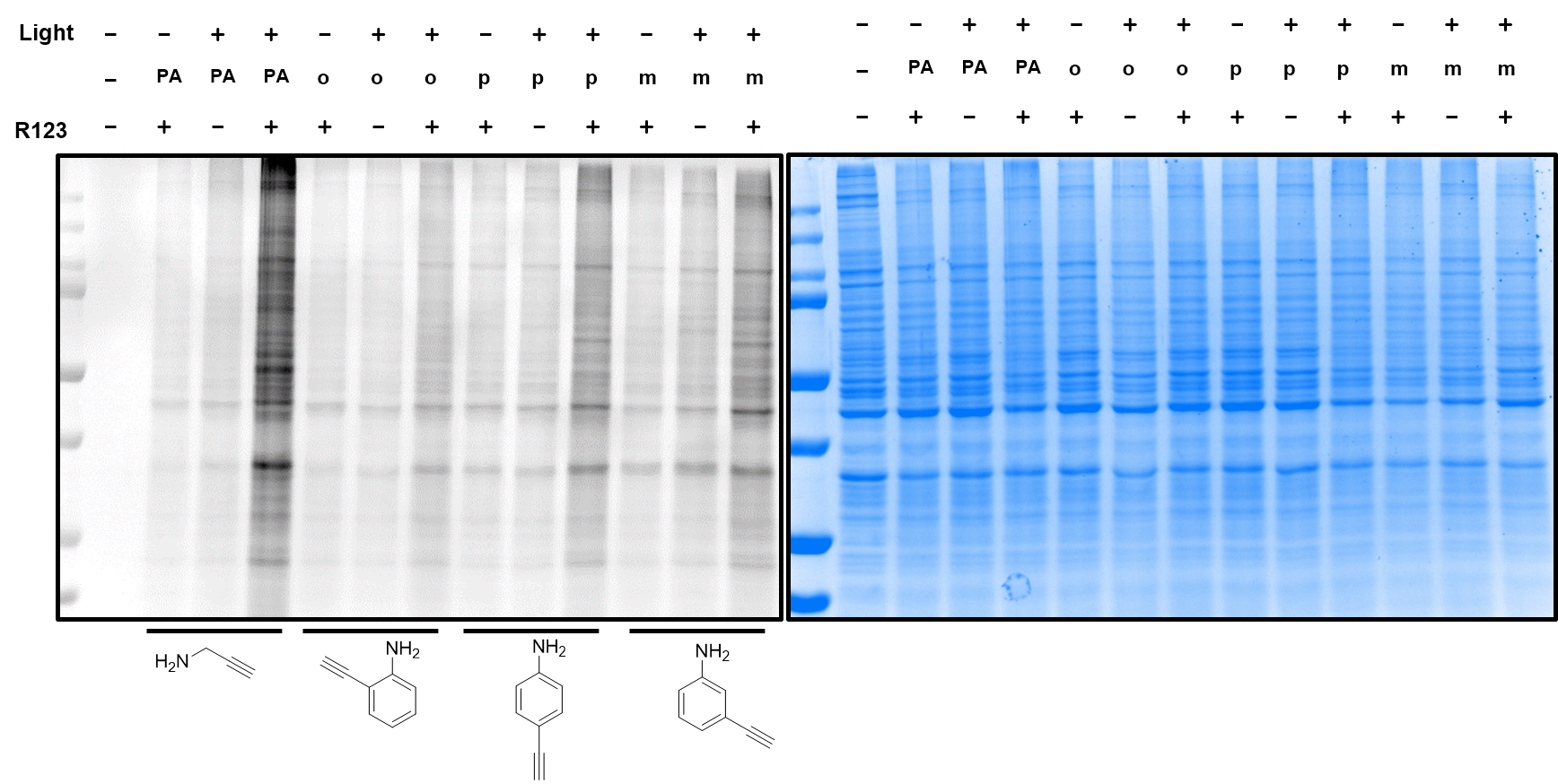

Fig. S6	Labeling efficiency of PA and o-, p-, m-aminophenylacetylenes mediated by R123 and light. HeLa Cells were stained with 10 μM R123 for 3h. Cells were treated with 5 mM AY or PA for extra 5 min. Cells were then illuminated for 5 min. Cell lysate were reacted with azido Cy5 for in gel fluorescence analysis.

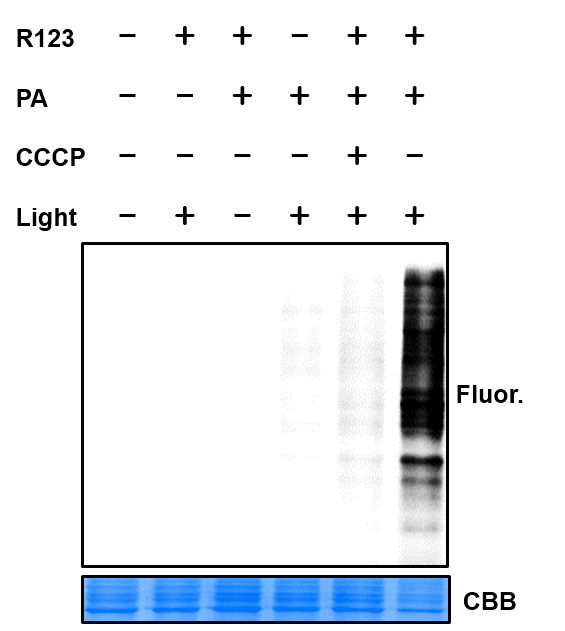

Fig. S7	In gel fluorescence analysis of mitochondrial proteins labeling with R123 and PA based LIMPLA2.

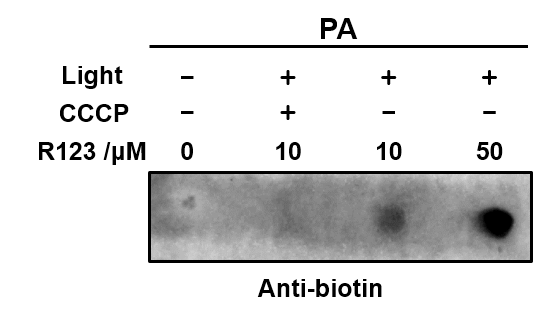

Fig. S8	Dot blot analysis of RNA labeling by LIMPLA2. HeLa cells were treated with 10 or 50 μM R123 (or supplemented 10 μM CCCP) for 2 h, cultured with 5 mM PA for 5 min. RNA were isolated from these cells and click labeled with azido biotin. Purified RNA were analyzed by dot blot analysis.

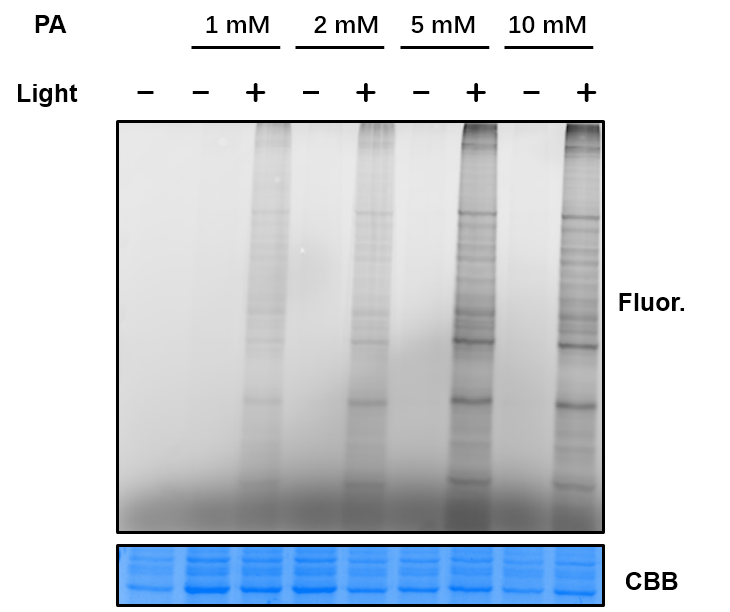

Fig. S9	PA concentration dependent labeling mediated by mito405 and PA based LIMPLA2.

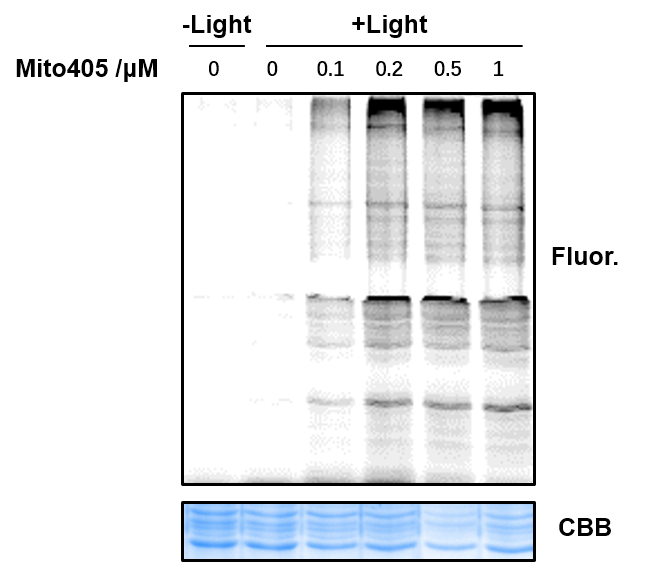

Fig. S10	Mito405 concentration dependent labeling mediated by mito405 and PA based LIMPLA2.

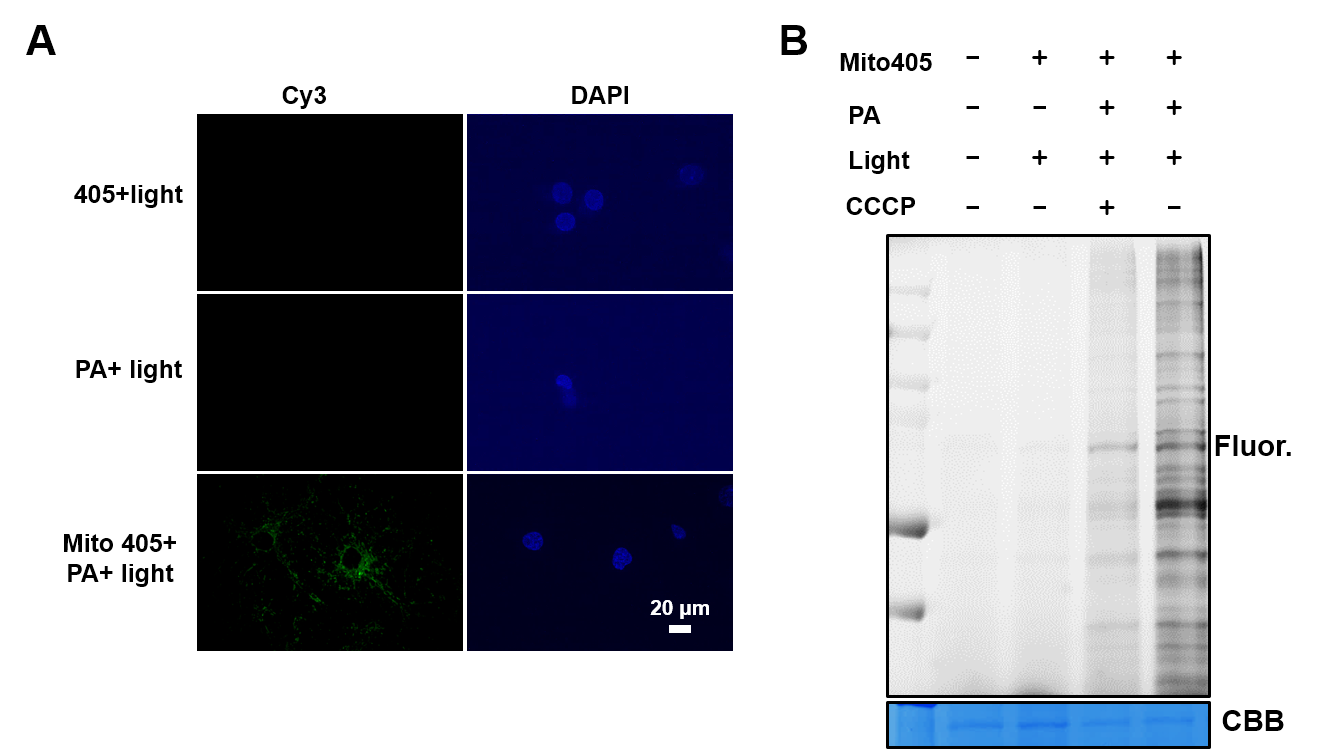

Fig. S11	Cortical neuron cells labeling with LIMPLA2. (A) Cortical neuron cells were stained with 0.1 μM mito405 for 2h, incubated with 5 mM PA for 5 min and illuminated for 5 min. Cells were clicked labeled with azido-Cy3 and analyzed with in situ fluorescent imaging (B) In gel fluorescence scanning analysis of labeling.

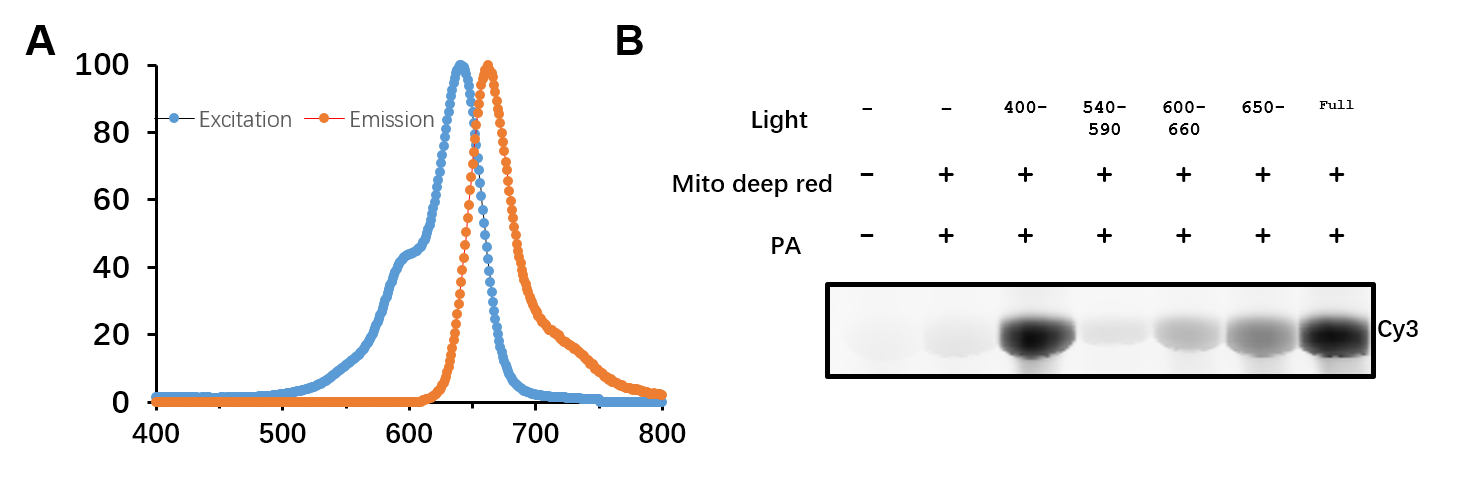

Fig. S12	BSA labeling mediated by Mito tracker deep red with various wavelengths of light. (A) Excitation and emission spectrum of Mitotracker deep red. (B) In gel fluorescence scanning analysis of protein labeling mediated with various wavelengths of light. 1mg/mL BSA mixed with 1 μM Mitotracker deep red and 1mM PA were illuminated with xenon lamp or xenon lamp with 400nm-, 540-590nm, 600-660nm filter for 2min. After protein precipitation, BSA were click labeled with azido Cy3.

Alk-R123:

^1^H NMR (400 MHz, Methanol-d4) δ 8.30 (dd, J = 7.8, 1.4 Hz, 1H), 7.84 (dtd, J = 25.5, 7.5, 1.4 Hz, 2H), 7.44 (dd, J = 7.5, 1.4 Hz, 1H), 7.03 (d, J = 8.9 Hz, 2H), 6.88 – 6.73 (m, 4H), 4.60 (d, J = 2.5 Hz, 2H), 2.77 (t, J = 2.5 Hz, 1H). HRMS (ESI): **369.12327.**

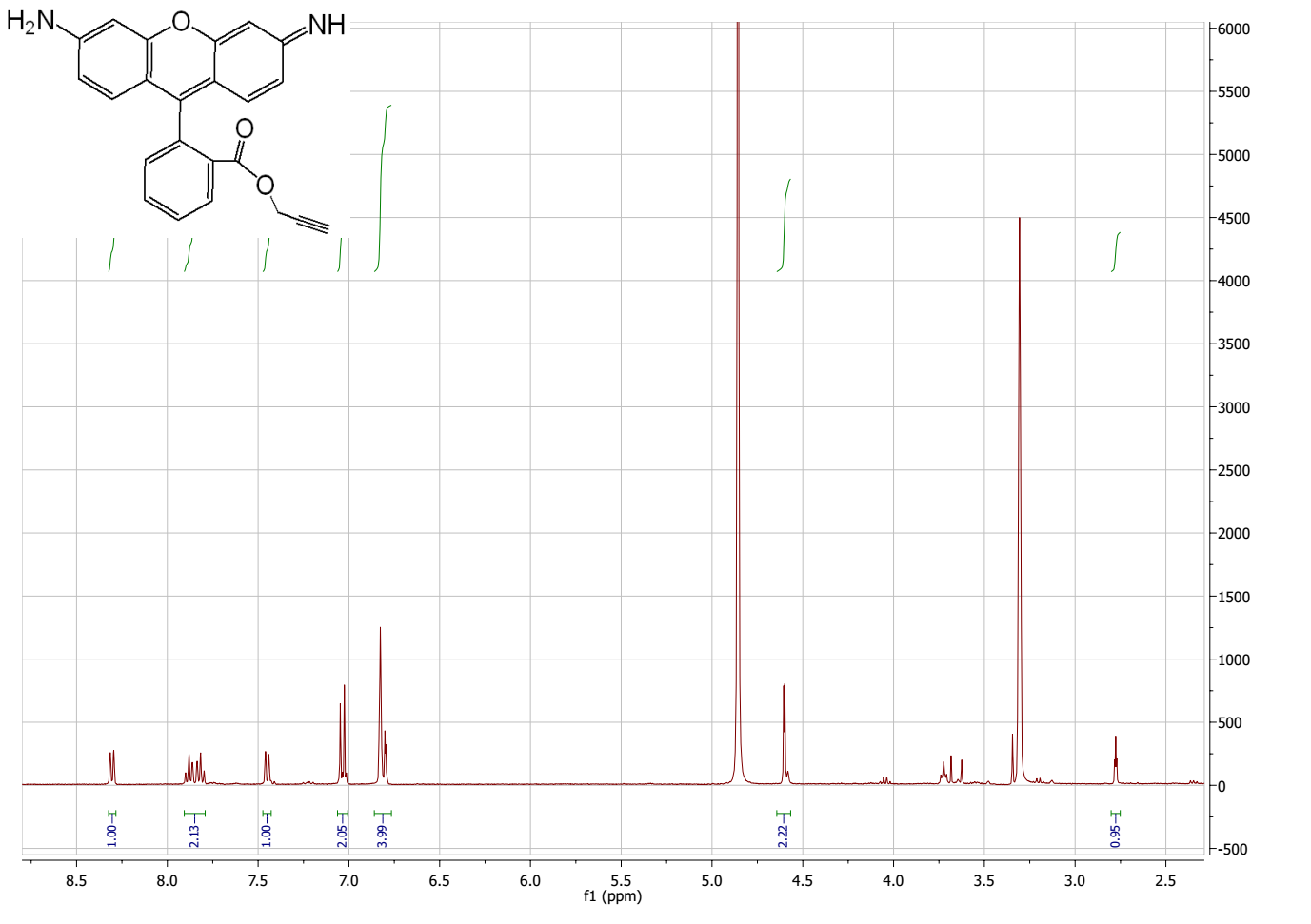

Az-R123:

^1^H NMR (400 MHz, Methanol-d4) δ 8.34 (dd, J = 7.8, 1.4 Hz, 1H), 7.86 (dtd, J = 23.8, 7.6, 1.4 Hz, 2H), 7.46 (dd, J = 7.4, 1.4 Hz, 1H), 7.07 (d, J = 9.0 Hz, 2H), 6.92 – 6.74 (m, 4H), 4.17 – 4.06 (m, 2H), 3.31 – 3.23 (m, 2H). HRMS (ESI): **400.14145.**
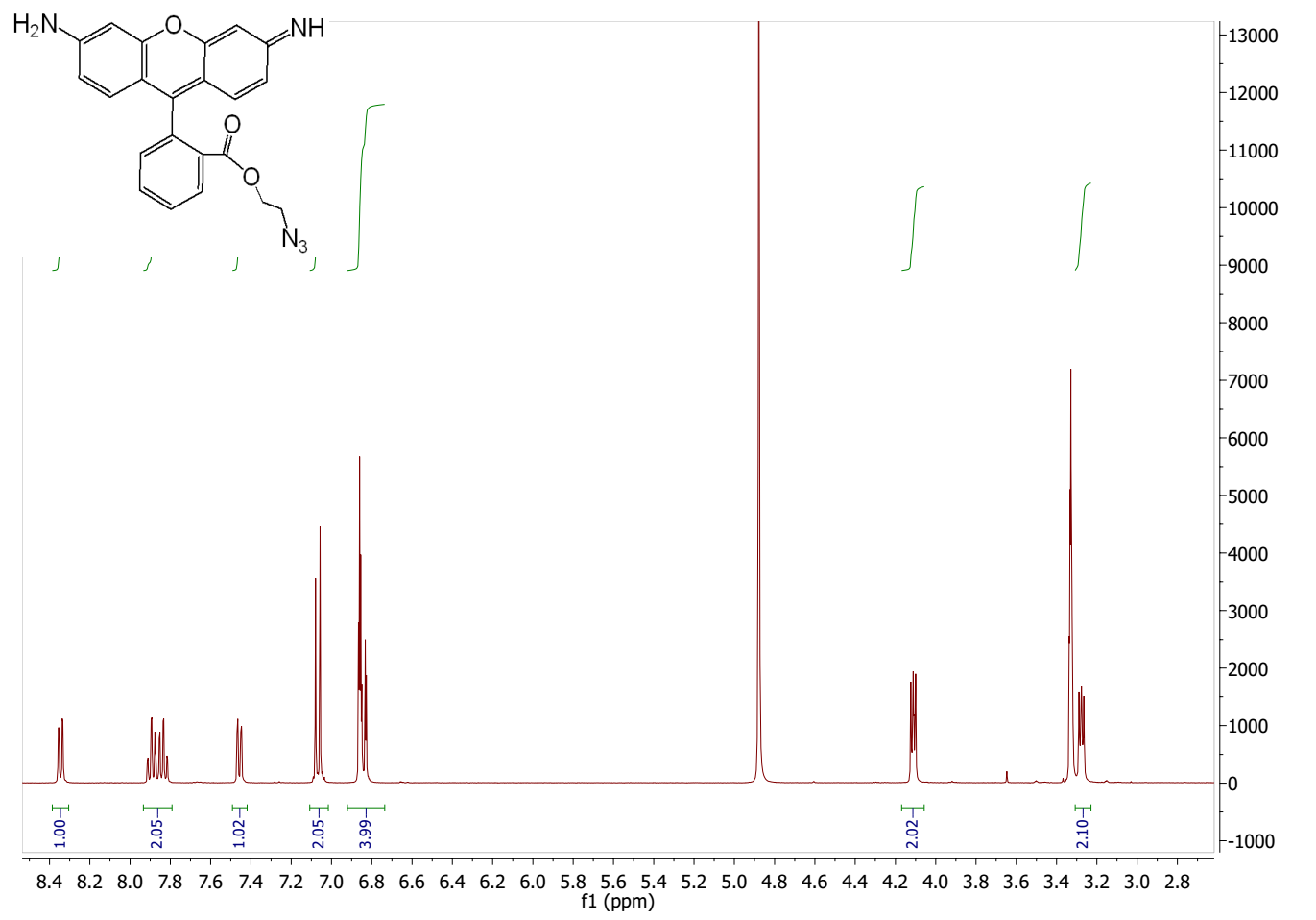
